# Insights into the mechanisms of LOV domain color tuning from a set of high-resolution X-ray structures

**DOI:** 10.1101/2021.02.05.429969

**Authors:** Alina Remeeva, Vera V. Nazarenko, Kirill Kovalev, Ivan Goncharov, Anna Yudenko, Roman Astashkin, Valentin Gordeliy, Ivan Gushchin

## Abstract

Light-oxygen-voltage (LOV) domains are widespread photosensory modules that can be used in fluorescence microscopy, optogenetics and controlled production of reactive oxygen species. All of the currently known LOV domains have absorption maxima in the range of ∼440 to ∼450 nm, and it is not clear whether they can be shifted significantly using mutations. Here, we have generated a panel of LOV domain variants by mutating the key chromophore-proximal glutamine amino acid of a thermostable flavin based fluorescent protein CagFbFP (Gln148) to asparagine, aspartate, glutamate, histidine, lysine and arginine. Absorption spectra of all of the mutants are blue-shifted, with the maximal shift of 8 nm observed for the Q148H variant. While CagFbFP and its Q148N/D/E variants are not sensitive to pH, Q148H/K/R reveal a moderate red shift induced by acidic pH. To gain further insight, we determined high resolution crystal structures of all of the mutants studied at the resolutions from 1.07 Å for Q148D to 1.63 Å for Q148R. Whereas in some of the variants, the amino acid 148 remains in the vicinity of the flavin, in Q148K, Q148R and partially Q148D, the C-terminus of the protein unlatches and the side chain of the residue 148 is reoriented away from the chromophore. Our results explain the absence of color shifts from replacing Gln148 with charged amino acids and pave the way for rational design of color-shifted flavin based fluorescent proteins.

## Introduction

Light-oxygen-voltage (LOV) domains are widespread photosensory modules found in proteins from organisms belonging to all three kingdoms of life^1,2^. They bind different flavins such as flavin mononucleotide (FMN) or flavin adenine dinucleotide (FAD) and respond to illumination with blue light^1,3,4^. At the moment, more than 6000 LOV sequences may be found in genomic databases^2^, and many of them have been characterized in detail. They generate considerable interest since they regulate a wide variety of processes such as phototropism in plants, circadian expression in fungi and lifestyle and virulence in bacteria^1,5^. Also, LOV domains have been engineered for applications in fluorescence microscopy^6–11^, optogenetics^12,13^ and controlled production of reactive oxygen species^14,15^. Among the major advantages of using LOV domains are their small size (around 110 amino acids) and independence of maturation on molecular oxygen or any other exogenous molecules.

The optical properties of the LOV domains are dictated by the flavin chromophores. The proteins have characteristic absorption spectra with the maxima at 370 and 450 nm, respectively. Absorption of a photon by the flavin launches a cascade of photochemical reactions called photocycle, where two widely conserved nearby amino acids, cysteine and glutamine, play a major role^1,3,4^. Cysteine’s sulfur forms a covalent bond with flavin’s C4a atom in the early steps of the photocycle, which leads to reorganization of the flavin’s hydrogen bonding pattern and, in particular, to change in the conformation of the glutamine^1,3,4^. Reorientation of the glutamine side chain further relays the photosignal^1,3,4^. Eventually, the cysteine-flavin bond breaks, and the protein returns thermally to the initial state^1,3,4^.

The role of different amino acids surrounding flavins in the photocycle of LOV domains has been extensively studied. Replacing the conserved cysteine with another sulfur-containing amino acid, methionine, leads to irreversible light-induced formation of the methionine-FMN adduct^16^. Replacing the cysteine with non-reactive amino acids abolishes the adduct formation^17,18^ and enhances the fluorescent properties of the protein^6,7^, because the adducted form is non-fluorescent. Yet, some LOV domains lacking the conserved cysteine are still able to generate a response to illumination^19^.

While the cysteine-FMN interaction is probably the most important one for the photocycle in most of the LOV domains, other amino acids regulate the duration of the photocycle to a high degree. The photocycles can last from seconds to days, and can be effectively regulated by mutations^20–23^.

Early mechanistic studies have also highlighted the role of a conserved glutamine, situated closely to FMN and the cysteine, as another key amino acid for signal transduction^24–27^. Replacement of the glutamine with a leucine has been used to remove the hydrogen bond with FMN, and impairs the signal transduction^22,27–30^. On the contrary, replacement with asparagine locks the LOV domain in an illuminated-like state and has been used as its mimic^30–33^. Finally, replacements of the glutamine with an aspartate or histidine have profound effects on the LOV domain photochemistry^22,34^.

Yet, none of the tested mutations had a profound effect on the absorption spectra of the LOV domains. While the absorption in the UVA region (320-400 nm) is affected by the amino acids surrounding the flavin’s 7a-methyl group^35^, the visible light spectra of the mutants are usually only slightly blue-shifted, with the maximum shift of 10 nm for Gln→Leu replacements^22,29,30,36^.

This highlights an intriguing question: can the visible light absorption of LOV domains be color-tuned at all? The native LOV domain-containing proteins seem all to have very similar spectra, with the absorption maxima of different proteins all being within several nanometers of each other (depending slightly on the equipment used in experiments), whereas other photoproteins display a remarkable variety of colors. For example, animal and microbial rhodopsins may have absorption maxima in essentially the whole range of the visible light from at least 400 to 700 nm, without any covalent modification of the retinal chromophore^37,38^. Beta-barrel fluorescent proteins such as the green fluorescent protein (GFP), or related blue, cyan, yellow and red FPs, have absorption maxima from ∼350 to ∼650 nm, although in this case the chromophores do differ^39–41^.

The common rationales in the studies of color tuning are that the spectra may be affected either by steric interactions (changing the geometry of the chromophore, which is not always possible), or electronic interactions (such as π-stacking, hydrogen bonding, or electrostatic)^42^. Since the isoalloxazine moiety in FMN and FAD is formed by three conjugated unsaturated heterocycles, substantially changing its geometry does not seem to be a viable idea. On the other hand, affecting its spectral properties by changing the charge distributions of the surrounding amino acids seems possible. In 2015, Khrenova *et al*. suggested that replacing the conserved glutamine 489 with a lysine in a widely used LOV-based fluorescent protein iLOV may result in a ∼50 nm red shift^43^. However, the experiments and calculations by Davari *et al*. showed that the Q489K mutant of iLOV is unexpectedly blue-shifted, and that the lysine’s ε-amino group is unlikely to remain in the close vicinity of the FMN’s isoalloxazine ring^44^. Consequently, Khrenova *et al*. suggested, based on simulations, that introducing a second substitution may fix the lysine side chain and result in a red shift^45^. Overall, electrostatic spectral tuning maps can be used to estimate the effects of the redistribution of charges around the chromophore on its spectra, in particularly for FMN-binding proteins^42^.

An alternative route to introducing mutations is modification of the chromophore. Replacement of FMN with roseoflavin strongly alters the LOV domain spectra and shifts the absorption maximum to 485 nm^46^. Replacement of FMN with 7-methyl-8-chloro-riboflavin has a similar effect^47^, whereas replacement with 7-bromo-8-methyl-riboflavin does not have a substantial effect, and replacement with lumichrome causes a blue shift of the absorption maximum to 421 nm^47^. Replacement of FMN with 5-deazaflavin mononucleotide similarly results in blue shifts to 423 nm^48^ or even ∼400 nm^49^, whereas 1-deazaflavin mononucleotide produces a remarkable red shift to ∼530 nm^49^. Finally, the absorption maxima of a LOV domain 8-bromo- and 8-trifluoromethyl-substituted flavins are relatively not shifted^50^. Although not yet confirmed experimentally, computational studies indicate that thioflavins or fluorinated flavins may also be used^51,52^, and that red shifts of up to 100 nm may also be attainable by combining modified chromophores with protein mutations^53^. Still, we must note that using the modified chromophores for applications of LOV domains does not seem to be particularly practical. While the natural chromophores such as FMN or FAD are ubiquitous in most living cells, the modified chromophores will have to be synthesized and supplemented exogenously for *in vivo* applications; they may be toxic for the cells, or their binding may be weaker compared to omnipresent natural flavins.

Therefore, a search for mutations that may shift the spectra of LOV domains, both to understand the color tuning mechanisms of flavins, and for applications, is fully justified. Among the amino acids surrounding the chromophore in LOVs, most are tightly constrained by their neighbors and/or the flavin, so that drastic mutations are very likely to result in loss of binding or misfolding. The only residue shown to tolerate some mutations to polar and charged amino acids is the glutamine in the vicinity of flavin atoms N4 and O5, which is involved in signaling. Previously, mutations of this glutamine were suggested to confer large spectral shifts^43,45^. Consequently, in this work, we have generated a panel of variants of the thermostable LOV protein CagFbFP, where the conservative glutamine is substituted with different polar and charged amino acids (aspartate, glutamate, lysine, arginine, asparagine and histidine) that might strongly affect the electrostatic environment of the flavin and thus change the absorption spectrum significantly. We showed that the mutations are tolerated and the protein remains folded and binds flavin, measured the absorption and fluorescence spectra of the generated variants, and determined their high-resolution crystallographic structures that allow us to rationalize the effects of mutations.

## Materials and Methods

### Cloning, protein expression and purification

The nucleotide sequence encoding the original variant CagFbFP (amino acid sequence derived from the gene Cagg_3753, UniProt accession code B8GAY9, *Chloroflexus aggregans* strain MD-66 / DSM 9485, with the mutation C85A and C-terminal 6×His tag) was obtained as a synthetic gene from Evrogen (Russia) and introduced into the pET11 expression vector (Novagen, Merck, Germany) via NdeI and BamHI restriction sites as described previously^54^. Mutations Q148N, Q148D, Q148E, Q148H, Q148K and Q148R were introduced into the CagFbFP gene using polymerase chain reaction.

CagFbFP and its mutants were expressed in *Escherichia coli* strain C41 (DE3) as described previously^54^. All cultures were grown in ZYP-5052 autoinducing medium^55^ supplemented with 100 μg mL^-1^ ampicillin at 37 °C. Proteins were purified using Ni-NTA resin (Qiagen, Germany) on a gravity flow column followed by size-exclusion chromatography on a Superdex® 200 Increase 10/300 GL column (GE Healthcare Life Sciences, USA) in a buffer containing 10 mM NaCl and 10 mM sodium phosphate, pH 8.0. Protein-containing fractions were concentrated to 30 mg ml^-1^ for crystallization.

### Spectroscopic methods

Thermal stability of CagFbFP and all mutated variants were measured using the Rotor-Gene Q real-time PCR cycler (Qiagen, Germany) with fluorescence excitation at 470 nm and fluorescence detection at 510 nm. Absorption and fluorescence spectra were recorded using Synergy™ H4 Hybrid Microplate Reader (BioTek, USA). Emission was measured between 470 nm and 700 nm in increments of 1 nm, with excitation fixed at 450 nm. Excitation was measured between 250 nm and 500 nm in increments of 1 nm, the signal was detected at 510 nm.

### Crystallization

All of the crystals were obtained using the sitting drop vapor diffusion approach. Crystallization trials were set up using the NT8 robotic system (Formulatrix, USA). The crystals were grown at 22 °C. The drops contained 150 nL concentrated protein solution and 100 nL reservoir solution.

CagFbFP-Q148N was crystallized with 0.06 M Magnesium chloride hexahydrate, 0.06 M Calcium chloride dihydrate, 0.1M MES monohydrate, 0.1M Imidazole, pH 6.5, 20% v/v PEG 500 MME, 10% w/v PEG 20000.

Crystals of CagFbFP-Q148D and CagFbFP-Q148E were obtained with the following precipitant: 0.1 M DL-Glutamic acid monohydrate, 0.1 M DL-Alanine, 0.1 M Glycine, 0.1 M DL-Lysine monohydrochloride, 0.1 M DL-Serine, 0.1M MES monohydrate, 0.1M Imidazole, pH 6.5, 20% v/v PEG 500 MME, 10% w/v PEG 20000.

Crystals of CagFbFP-Q148H were obtained using 0.5 M Lithium sulfate monohydrate, 15 % w/v PEG 8000 as precipitant.

As described in the accompanying article^56^, CagFbFP-Q148K was crystallized using the precipitant solution containing 0.1 M MES monohydrate pH 6.5 and 12% w/v PEG 20000. Morpholine-free structures were obtained with 0.5 M Ammonium Sulfate, 1 M Lithium sulfate monohydrate, 0.1 M Sodium citrate tribasic dihydrate pH 5.6.

Crystals of CagFbFP-Q148R were obtained with the following precipitant: 0.12 M 1,6-Hexanediol, 0.12 M 1-Butanol, 0.12 M 1,2-Propanediol, 0.12 M 2-Propanol, 0.12 M 1,4-Butanediol, 0.12 M 1,3-Propanediol, 0.1M MES monohydrate, 0.1M Imidazole, pH 6.5, 20% v/v PEG 500 MME, 10% w/v PEG 20000 as precipitant.

To prevent drying and for cryoprotection, precipitant solutions with 20% glycerol were added to the crystallization drops during crystal harvesting. The crystals were harvested using micromounts and then flash-cooled and stored in liquid nitrogen.

### Acquisition and treatment of diffraction data

The diffraction data were collected at 100 K on the beamlines ID23-1, ID29 and ID30B at the European Synchrotron Radiation Facility (ESRF, Grenoble, France). Diffraction images were processed using XDS^57^. POINTLESS and AIMLESS^58^ were used to merge, scale and assess the quality of the data, as well as to convert intensities to structure factor amplitudes and to generate Free-R labels. As described in the accompanying article^56^, datasets 6YX6 and 6YXC revealed strong anisotropy and were processed using STARANISO^59^ using the following criterion for determining the diffraction-limit surface: <I_mean_/σI_mean_> of more than 2.

### Structure determination and refinement

All structures were solved using molecular replacement with MOLREP^60^. CagFbFP structure^54^ (PDB ID 6RHF) was used as a search model. The resulting solution was refined manually using Coot^61^ and REFMAC5^62^.

## Results

### General properties of the Q148X variants

Introduction of strongly charged amino acids at the chromophore-binding site may affect its binding, and also destabilize the protein overall. Therefore, we speculated that starting with an initially stable protein, which can also produce well-diffracting crystals, is desirable for this study, and chosen the recently identified thermostable LOV domain from *Chloroflexus aggregans*^54^ as a framework. We used the C85A mutant, dubbed CagFbFP (*Chloroflexus aggregans* flavin based fluorescent protein), which is fluorescent and does not undergo a photocycle under illumination. Below, the designation WT is used for the original CagFbFP variant.

All of the tested variants (Q148N, Q148D, Q148E, Q148H, Q148K and Q148R) were expressed in *E. coli* in the flavin-bound form and could be purified and crystallized. We tested the effects of the substitutions on thermal stability of CagFbFP by measuring the dependence of fluorescence on temperature (Figures 1 and S1, and Table 1) and found that all of the substitutions destabilized the protein. Similarly to CagFbFP, the variants had complex melting curves, often with two different melting transitions. Among the variants tested, Q148K is the most detrimental mutation, with the melting temperature reduced by ∼20 °C.

**Figure 1.**
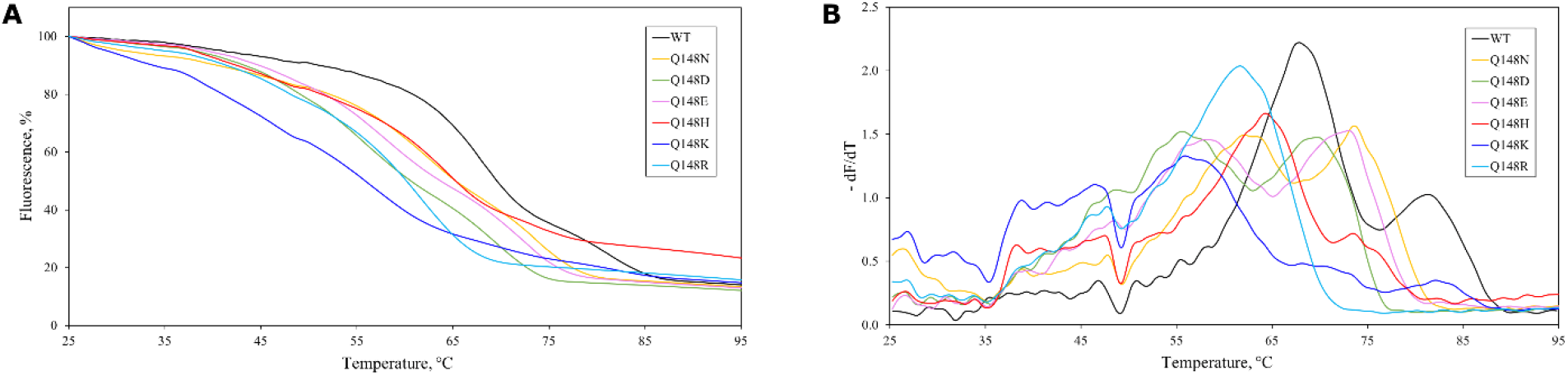
Dependence of CagFbFP variants’ fluorescence on temperature during heating. Temperature-induced decay of fluorescence. The values are normalized to 100% at 25 °C, the temperature in the beginning of the experiment. B) Derivatives of the fluorescence traces. Each experiment was conducted independently for five times, and the data were averaged for plotting. The troughs observed at 48 °C are due to the instrument and are observed in all samples. Characteristic unfolding temperatures are summarized in Table 1. Refolding data is shown in Figure S1.

**Table 1.** Properties of glutamine-substituted CagFbFP variants.

| Variant | T<sub>m1</sub>, °C | T<sub>m2</sub>, °C | T<sub>r</sub>, °C | λ<sub>ex</sub>, nm | λ<sub>em</sub>, nm |
| --- | --- | --- | --- | --- | --- |
| WT | 67.9 ± 0.3 | 81.3 ± 0.3 | 65.4 ± 0.3 | 450 | 498 |
| Q148N | 62.2 ± 1.0 | 73.6 ± 0.3 | 61.7 ± 0.3 | 447 | 496 |
| Q148D | 55.7 ± 0.3 | 69.0 ± 0.7 | 45.3 ± 0.7*** | 449 | 501 |
| Q148E | 58.3 ± 0.6 | 72.4 ± 0.9 | 49.2 ± 1.3 | 449 | 505 |
| Q148H | 64.3 ± 0.3 | 73.5 ± 0.2 | 65.4 ± 0.3 | 444 | 492 |
| Q148K | 56.0 ± 0.5 | n.d.** | 55.6 ± 1.9*** | 446 | 492 |
| Q148R | 61.6 ± 0.4 | n.d.** | 48.7 ± 2.1*** | 447 | 499 |

\* The errors are standard deviations of the peak positions in five melting experiments.
\*\* For these variants, presence and position of the second melting transition is not clear.
\*\*\* Only a small fraction is refolded, the temperature is not reliable.

### Spectroscopic characterization of the Q148X mutants

Having generated the Q148X mutants, we measured their absorption and fluorescence spectra (Figure 2 and Table 1). We note that positions of absorption and fluorescence excitation maxima are slightly different, with the former being blue-shifted by 2-3 nm compared to the latter.

**Figure 2.**
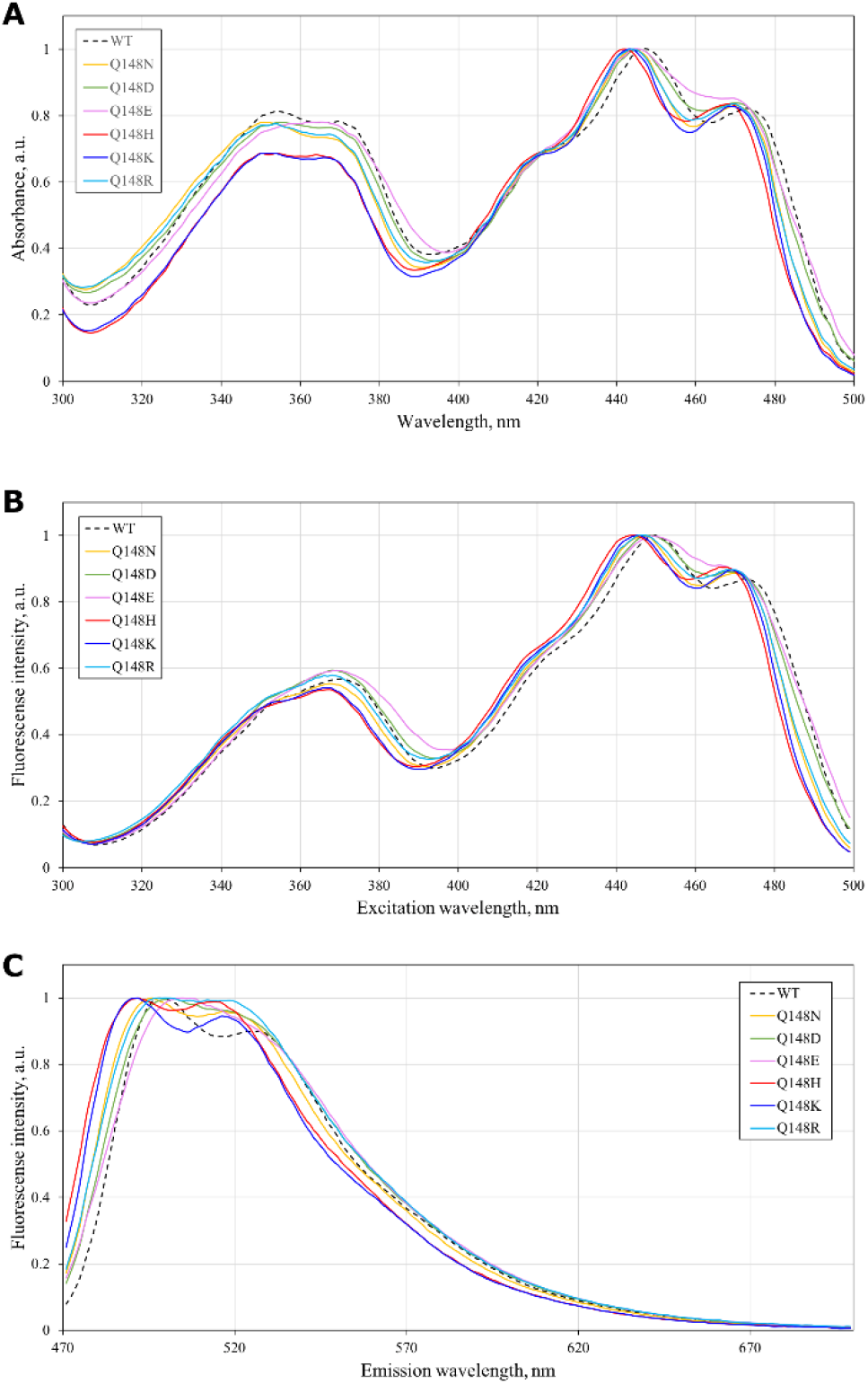
Spectroscopic properties of CagFbFP variants at pH 8. A) Absorption spectra. Fluorescence excitation spectra. C) Fluorescence emission spectra. The data are summarized in Table 1. Absorption and excitation data for the wavelength range 250-500 nm are presented in Supporting Figure S2.

Interestingly, none of the variants revealed a red shift in the absorption or fluorescence excitation spectrum; substitutions of the glutamine 148 with potentially positively charged amino acids resulted in the largest blue shifts, for as much as 8 nm for Q148H. The blue shift of the Q148K variant is smaller than that of the homologous iLOV Q489K mutant^44^. While the fluorescence emission maxima of the Q148D and Q148E variants are formally red-shifted, the spectra are overall widened and cannot be considered red-shifted.

### Structural analysis of the Q148X mutants

Following the spectroscopic characterization, we attempted crystallization of all of the generated variants. The original variant, CagFbFP, has been crystallized previously, and its structure has been determined at the resolution of 1.07 Å^54^. The new variants are also readily crystallizable, and X-ray structures were solved for all of them at the resolutions ranging from 1.07 Å for Q148D to 1.63 Å for Q148R. All of the crystals belonged to the same space group as CagFbFP crystals, P2_1_2_1_2. For Q148H and Q148K, crystals belonging to C2 and P2_1_, respectively, were also obtained. In comparison to the P2_1_2_1_2 CagFbFP crystals, P2_1_2_1_2 Q148K and Q148R crystals had slightly extended unit cell dimension a (∼57.5 Å instead of ∼53.5 Å). The structures have been deposited to the Protein Data Bank (PDB), and respective PDB identifiers are used for convenience below. Crystallographic data collection and refinement statistics are presented in Supplementary Table S1. Crystals of the Q148K variant in the space group P2_1_ contained two dimers in the asymmetric unit; all other crystals contained a single dimer in the asymmetric unit. In many cases this was helpful as more data could be obtained from a single crystal.

Since aspartate, glutamate, histidine, lysine and arginine side changes can be protonated or deprotonated depending on pH and their environment, we tested the effects of pH on the fluorescence spectra (Figures 3 and S3-S7). The parent protein, CagFbFP, didn’t reveal any differences in the spectra collected at different pH values, similarly to EcFbFP, which retained its fluorescence in a wide range of pH values^63^. Fluorescence spectra of the variants Q148N,D,E were also essentially identical at pH 3 and pH 8 (Figures S3-5). On the contrary, Q148H,K,R fluorescence spectra were slightly red-shifted at pH 3 compared to pH 8 (Figures 3 and S6-7). A possible explanation for this will be given below in the context of the structural data.

**Figure 3.**
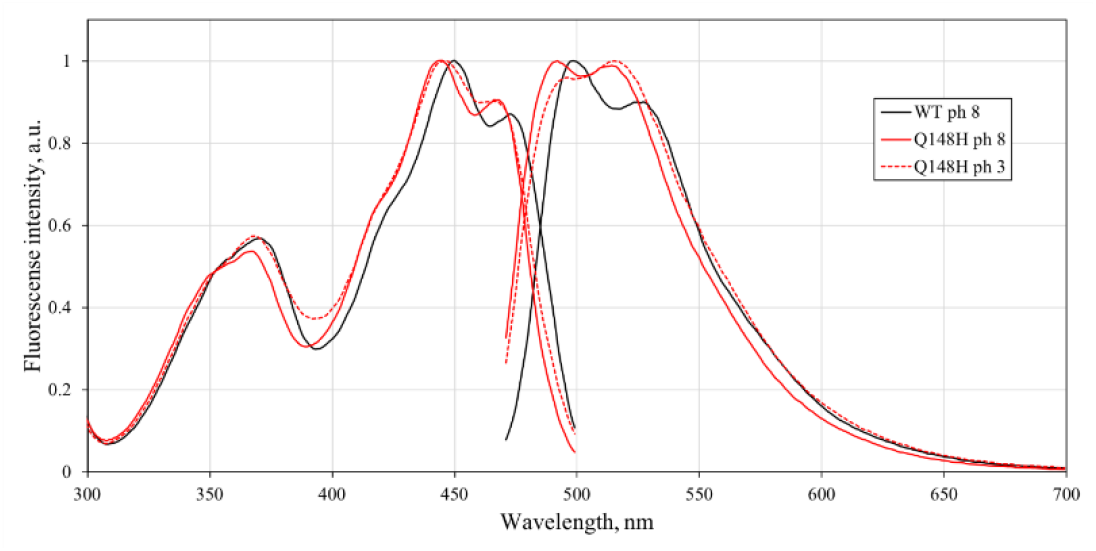
Dependence of CagFbFP Q148H fluorescence excitation and emission spectra on pH. All of the structures reveal the same fold of the proteins. Residues 48-148 are well resolved in all of the structures, and their backbones occupy essentially identical positions. The C-terminus, starting from the residue 149, has sometimes different conformations, as do the amino acids Asn127 and Asn/Asp/Glu/His/Lys/Arg148. The obtained structures are depicted in Figures 4 and 5, and exemplary 2F_o_-F_c_ and omit (polder^64^) maps may be found in Figure S8.

### Structure of CagFbFP

To put the new findings in context, we describe first the structure of the original variant, CagFbFP^54^ (Figure 4A). The chromophore FMN is coordinated by, among others, Asn117 and Asn127 side chains, and Gln148 occupies two alternative conformations, in both cases forming a hydrogen bond to the FMN’s atom O4. Whereas one of the conformations is similar to that observed in most other LOV domains, the second one is unusual and may be a consequence of steric interactions with the nearby Ile52 side chain. The asymmetric unit contains two CagFbFP molecules, which are almost identical in structure. Yet, there is a small difference: in the chain A, the C-terminus occupies a single (latched) conformation, and separates the Gln148-harboring pocket from the solvent. However, in the chain B, two alternative conformations of the C-terminus are observed. In the second one, Thr149 is displaced, and Asp150 is not very well resolved in the electron density maps.

**Figure 4.**
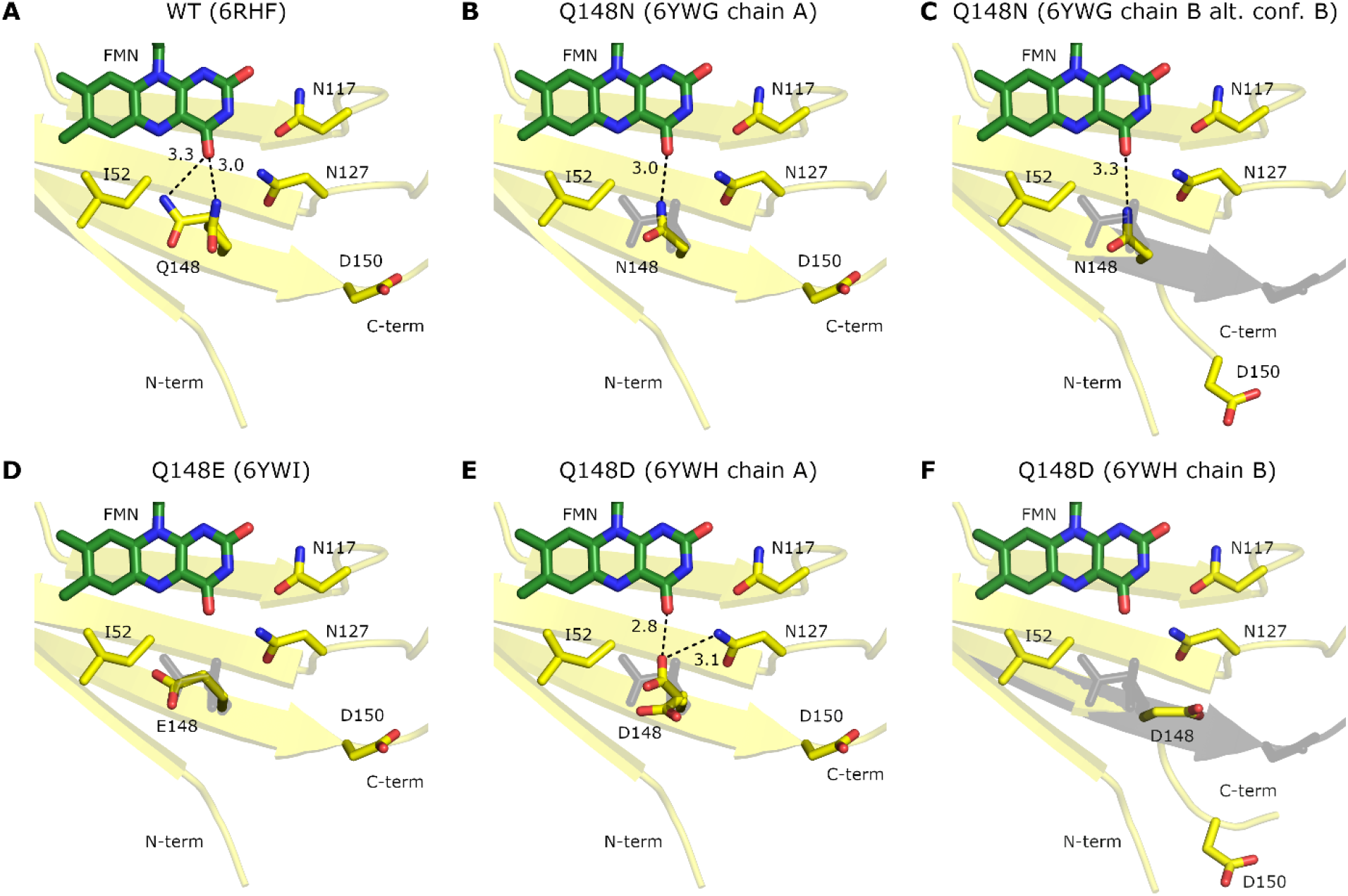
Structure of the flavin-binding pocket in CagFbFP and its Q148N, Q148E and Q148D mutants. Positions of CagFbFP’s Gln148 and C-terminus are shown in all panels in grey for reference. Putative hydrogen bonding distances are indicated in ångströms. A) Structure of CagFbFP that was determined previously^54^. Note that Gln148 was observed to occupy two alternative positions. B) Structure of the Q148N mutant, chain A. Asn148 forms a hydrogen bond with FMN. C-terminus occupies the same position as in the original variant. C) Structure of the Q148N mutant, chain A. Two alternative conformations of the C-terminus are observed: unlatched (shown) and latched, as in panel B (not shown). D) Structure of the Q148E mutant. No evident hydrogen bonding partners are observed for the Glu148 side chain. E) Structure of the Q148D mutant, chain A. Two alternative conformations of the Asp148 side chain are observed: in the first one, it forms a hydrogen bond with FMN, similarly to Asn148 in the Q148N mutant; in the second one, no evident hydrogen bonding partners are observed. F) Structure of the Q148D mutant, chain B. Asp148 side chain is reoriented away from FMN; C-terminus in unlatched.

### Structure of the Q148N variant

Similarly to glutamine, asparagine has a carboxamide side chain, and we begin our comparisons with the structure of the Q148N variant. In both chains of the dimer, Asn148 side chain occupies the same position (Figures 4B,C). Its atom N_δ2_ is within hydrogen bonding distance of the FMN’s O4. Whereas the protein’s C-terminus is in a single latched conformation in the chain A (Figure 4B), it is clearly in two conformations in the chain B: one is like that in the chain A, and another is different (Figure 4C). Overall, the structure is identical to that of the original variant CagFbFP in all regards except for the position of the residue 148 side chain. Consequently, it is not clear how the mutation may convert the protein into illuminated-like state, as it does in *Avena sativa* phototropin 1 LOV2^30^; perhaps, it changes the dynamics of the proteins in solution.

### Structure of the Q148E variant

As opposed to glutamine, glutamate has a carboxylic acid side chain that can become negatively charged if deprotonated. Surprisingly, we observe that Glu148 side chain occupies almost the same position in the Q148E variant as Gln148 does in the WT CagFbFP, except that it loses the hydrogen bond to FMN (Figure 4D). The distances from the O_ε1_ and O_ε1_ carboxyl atoms to FMN’s O4 are 4.3 and 5.1 Å, respectively, and there are no evident hydrogen bonding partners in the vicinity of the side chain. In both chains, the C-termini are clearly in the same single latched conformations. In the chain A, residues 149-155, including His154 and His155 of 6×His purification tag, are well resolved.

### Structure of the Q148D variant

Aspartate has a shorter side chain compared to glutamate, and this might be the reason for the observed differences in the structures (Figure 4E,F). Asp148 and C-terminus occupy different positions in the two protein chains in the asymmetric unit. In the chain A (Figure 4E), the C-terminus is latched, and positions of residues 149-155, including His154 and His155 of 6×His purification tag are resolved similarly to those in the Q148E variant. Asp148 occupies two alternative conformations: one within hydrogen bonding distance of the FMN’s O4 and Asn127’s N_δ2_, and another one rotated and without any evident hydrogen bonding partners in the vicinity. The hydrogen-bonded conformation is similar to that observed by Kopka *et al*. in a simulation of the Q489D variant of iLOV with protonated D489^34^. In the chain B, surprisingly, the C-terminus is fully unlatched, and Asp148 side chain is oriented away from FMN (Figure 4F), as observed in a simulation of the Q489D variant of iLOV with deprotonated D489^34^.

### Structure of the Q148K variant

As opposed to the previously discussed amino acids, lysine has an amino group in its side chain. It is positively charged and protonated when exposed to the solvent at physiological conditions, but its pK_a_ may be as low as 5.3^65^ when it is confined within a protein (as observed, for example, in some microbial rhodopsins^66^). Previously, both Khrenova *et al*.^43^ and Davari *et al*.^44^ proposed that the homologous Lys489 in the Q489K mutant of iLOV is protonated, and its ε-amino group remains within 6 Å of the FMN’s O4.

As described in the accompanying article^56^, we were able to obtain three types of Q148K variant crystals, all using the precipitant solutions with pH of 6.5. Unexpectedly, and contrary to the results of previous simulations^43,44^, in all of the obtained models the C-terminus of the protein is clearly unlatched, and the side chain of Lys148 is extended away from FMN (Figure 5A,B,C).

**Figure 5.**
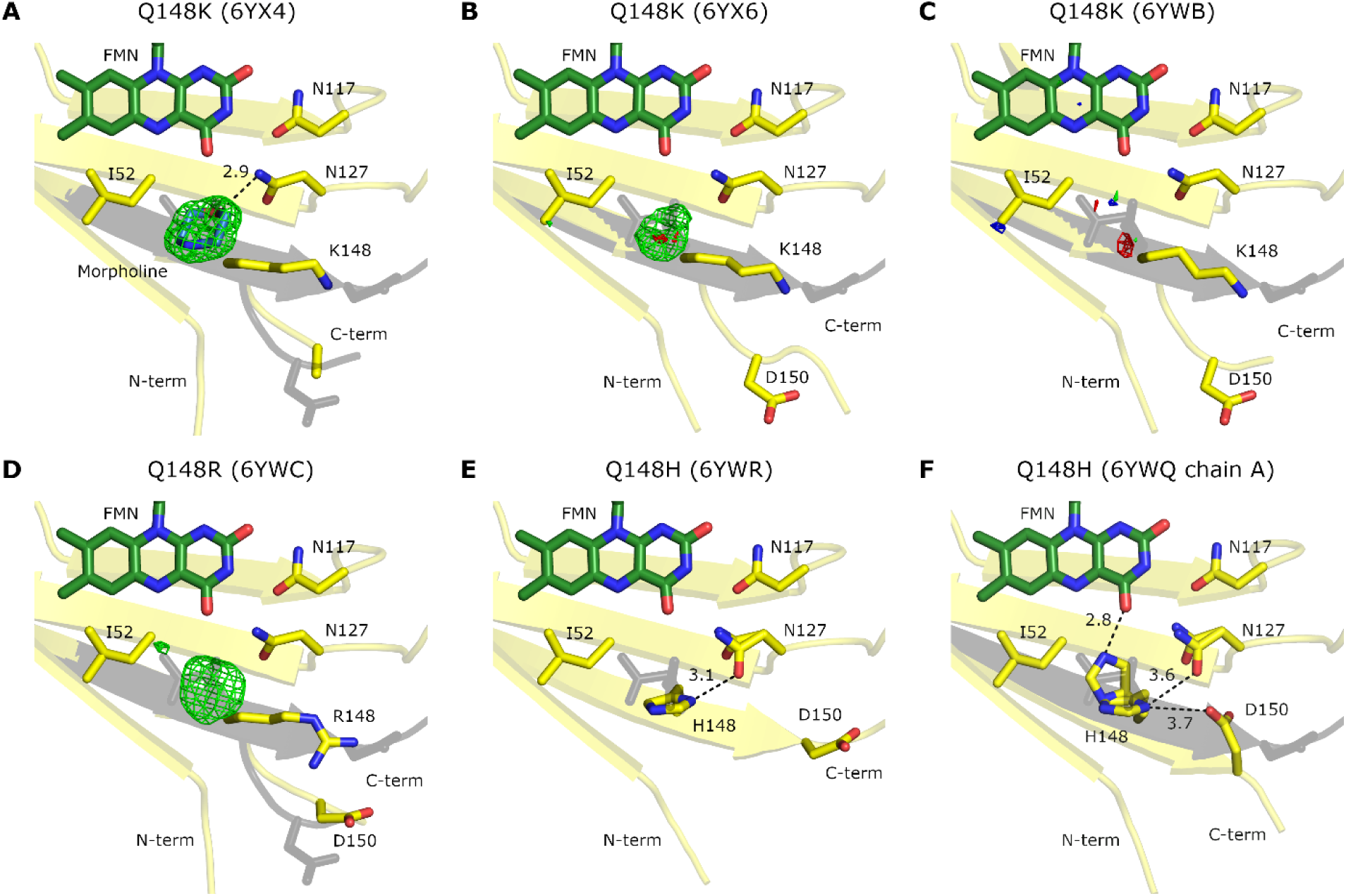
Structure of the flavin-binding pocket in Q148K, Q148R and Q148H mutants of CagFbFP. Positions of CagFbFP’s Gln148 and latched and unlatched C-terminus are shown in in grey for reference. A) Structure of the Q148K mutant with a bound morpholine molecule (blue). Polder omit map^64^ is contoured at the level of 4 × r.m.s. (green). Lys148 side chain is reoriented away from FMN; C-terminus in unlatched, but differently from what is observed for Q148N and Q148D mutants. Position of Asp150 side chain cannot be determined from the available electron densities. B) Structure of the Q148K mutant without morpholine (space group P2_1_2_1_2). Lys148 side chain is reoriented away from FMN; C-terminus in unlatched. Residual polder omit densities (green for chain A and red for chain B, contoured at the level of 3.5 × r.m.s.) are observed near FMN. C) Structure of the Q148K mutant without morpholine (space group P2_1_2_1_2). Residual polder omit densities (green for chain A, red for chain B, blue for chain D, contoured at the level of 3.5 × r.m.s.) are observed near FMN. Lys148 side chain is reoriented away from FMN; C-terminus in unlatched. D) Structure of the Q148R mutant. Polder omit map^64^ is contoured at the level of 3 × r.m.s. (green). Arg148 side chain is reoriented away from FMN; C-terminus in unlatched, but differently from what is observed for Q148N and Q148D mutants. E) Structure of the Q148H mutant (space group C2). Two alternative conformations for Asn127 are observed: similar to the one observed in other variants, and a new one hydrogen-bonded to the N_δ1_ atom of His148. F) Structure of the Q148H mutant (space group P2_1_2_1_2). Two alternative conformations for Asn127 are observed. Two alternative conformations for His148 are also observed: similar to the one observed in the C2 crystals, and one where His148 is hydrogen-bonded to FMN. Asp150 is reoriented towards His148.

Surprisingly, the highest-resolution structure (PDB ID 6YX4, determined at the resolution of 1.36 Å), revealed a six-membered ring in a chair conformation close to FMN (Figure 5A). No such molecule has been added to the sample during purification or crystallization. Yet, we noted that the precipitant solution contained 2-(N-morpholino)ethanesulfonic acid (MES), and morpholine moiety fitted perfectly in the density near FMN with the occupancies of 0.7 and 0.5 for the protein chains A and B, respectively. Thus, we assume that MES was degraded to morpholine during storage of the precipitant solution or during crystallization, and the morpholine bound to the pocket in CagFbFP. Similar presumable degradation of a related buffer component 3-(N-morpholino)propanesulfonic acid (MOPS) and binding of morpholine to the protein was observed by Sooriyaarachchi *et al*. during crystallization of *Plasmodium falciparum* 1-deoxy-d-xylulose-5-phosphate deductoisomerase^67^ (PDB ID 5JO0).

Although we did not observe any influence of morpholine on the Q148K spectra or thermal stability, it could not be excluded that it affected the conformation of the protein. Consequently, we crystallized the protein using another precipitant solution in two crystal forms^68^. P2_1_2_1_2 crystals (PDB ID 6YX6) diffracted anisotropically, and we applied ellipsoidal truncation using the resolution cut-offs of 1.5 and 2.05 Å. Chain A revealed some residual density in the vicinity of FMN observed at the level of the polder maps of 3.5 × r.m.s. (Figure 5B, green) that couldn’t be ascribed to any specific molecule. Chain B revealed only a very weak density at the same level (Figure 5B, red). P2_1_ crystals (PDB ID 6YXB) produced almost isotropic diffraction patterns and contained two protein dimers within the asymmetric unit. Chains A, B and D contained very weak densities at the same position (Figure 5C, green, red and blue), whereas chain C didn’t have any. The densities are likely due to disordered water molecules present in the pockets of some of the proteins in the crystal. Therefore, we conclude that irrespective of the buffer, the C-terminus of the Q148K variant is unlatched and Lys148 occupies the outermost position. These findings are corroborated by the structure of the Q489K variant of iLOV described in the accompanying article^56^.

### Structure of the Q148R variant

Arginine has a guanidinium group in its side chain that is also normally protonated and has an even higher pK_a_ in solution (∼12.5) compared to lysine (∼10.5). Similarly to Lys148 in the Q148K variant, Arg148 in Q148R occupies the outermost position (Figure 5D). There are two density blobs in the vicinity of FMN, a smaller one within hydrogen-bonding distance of N5 and O4 atoms, and a bigger one at the position of morpholine and other densities in the Q148K variant. The smaller density likely reflects partial occupancy of a water molecule, whereas the bigger one cannot be ascribed to any particular molecules in the precipitant solution. The C-terminus is in an intermediate conformation similar to that of the morpholine-bound Q148K variant. Interestingly, Asp150 side chain is nearby and may interact with the Arg148 side chain (Figure 5D).

### Structure of the Q148H variant

Histidine is the most difficult side chain for analysis and modeling since it has several protonation states, tautomeric forms, and in solution its pK_a_ is around 6.5. We obtained two crystal forms of Q148H, belonging to space groups P2_1_2_1_2 (PDB ID 6YWQ) and C2 (PDB ID 6YWR). In the first structure (PDB ID 6YWQ), in both chains, the C-terminus is in the latched conformation, and His148 side chain does not contact FMN (Figure 5E). Unlike in any of the previously discussed structures, Asn127 occupies two alternative conformations; in the new one it is hydrogen-bonded to His148 (Figure 5E). In the second structure (PDB ID 6YWR), the chain A adopts the same conformation, with possibly a weakly occupied second conformation of His148 attained via rotation around the C_β_-C_γ_ torsion. However, chain B has a number of unique features: besides the two alternative conformations for Asn127, two conformations are also observed for His148 (the new one is hydrogen-bonded to FMN), and Asp150 is reoriented towards His148 (Figure 5F).

## Discussion

Herein, we have generated and characterized a panel of 6 variants of a thermostable LOV domain CagFbFP, where the conservative glutamine amino acid has been replaced by asparagine, glutamate, aspartate, histidine, lysine or arginine. All of the tested mutations were destabilizing and affected the melting and refolding temperatures strongly (Table 1), which highlights the importance of starting with an initially stable protein^69^. Surprisingly, none of the mutations produced a red shift in the absorbance or fluorescence excitation spectra, and the observed blue shifts were smaller than those resulting from Gln→Leu replacement in other proteins^36^. Thus, significant color tuning would probably require two or more simultaneous mutations^45,56^.

We also determined 9 high-resolution crystal structures of the mutants (Table S1). The structures highlight the overall robustness of the LOV domain fold, and the conformations of various amino acids at the position 148. In the original version CagFbFP, residue 148 adopts two conformations: one is classical and observed in many other LOV domain structures, and the other is perpendicular to it^54^. This second conformation was implicated in the signaling mechanism in ZEITLUPE^70^ (dubbed the “exposed” conformation) and in *Bacillus subtilis* YtvA^71^. Interestingly, in the mutated variants, the side chains are in most cases observed in a single conformation (Figures 4 and 5). In particular, the structures reveal that the potentially strongly charged amino acids (Lys, Arg, and partially Asp), which might have affected the spectroscopic properties of the flavin chromophore, are in fact expulsed from the protein core (Figures 4 and 5). This explains the seeming contradiction between the notion that the strong positive charges in the region of the flavin’s N5 and O4 atoms should result in a strong red shift^42–44^, and the experimental data revealing modest blue shifts (reported by Davari *et al*.^44^, here, and also in the accompanying article^56^). Potentially negatively charged amino acids also do not produce a substantial blue shift, contrary to what might have been expected from the electrostatic tuning maps^42,72^. Evidently, Glu148 and Asp148 either do not interact with FMN, or interact with it in the protonated form (hydrogen-bonded conformation in Figure 4E). These findings underscore the importance of using experimental structural data for prediction of properties of photoactive proteins^73^.

Our data also shows that Q148H, K and R mutants are mildly pH-sensitive, revealing red shifts at lower pH (Figures 3 and S6-7). This sensitivity comes from the mutations, because CagFbFP and the spectra of the variants Q148N, D and E do not depend on pH (Figures S3-5). At the same time, Lys148 and Arg148 in Q148K and Q148R are highly likely to be protonated and charged already at physiological pH and thus should not be affected by lower pH values. We believe that lower pH values may result in neutralization of Asp150, which might serve as a counterion for Lys148 or Arg148 (Figure 5A-D), and consequently in a higher probability of lysine/arginine side chain coming in the vicinity of FMN and affecting its spectra.

As can be seen from the structures, the reorientation of the residue 148 side chain away from the chromophore in Q148K, Q148R and partially Q148D variants is possible due to unlatching of the C-terminus of the LOV domain. Thus, it might be possible to keep the residue 148 side chain nearby the chromophore either by creating favorable interactions in the pocket, as suggested by Khrenova *et al*.^45^, or by stabilizing the C-terminus in the latched conformation. The former is demonstrated in the accompanying article^56^. On the other hand, unlatching and opening of FMN-proximal pocket presents an interesting possibility of influencing the LOV domain spectra by small molecules, or even designing LOV-based fluorescent reporters that change their properties upon binding of a particular molecule.

Finally, our study highlights CagFbFP as a useful framework for engineering of new LOV domain variants. It clearly tolerates drastic alterations in the FMN-binding pocket, and even then, it still remains relatively thermostable. CagFbFP can be split for detection of protein-protein interactions^74^. It can be further thermostabilized by replacing Ala95 with a proline^75^ for testing even more destabilizing alterations. Finally, CagFbFP can be crystallized and produce high-resolution structures on par with the best resolution structures available for other LOV proteins (1.17 Å for miniSOG^15^ and 1.2 Å for iLOV^76^).

## Supporting Information

Supporting information (Supporting Table S1 and Supporting Figures S1-S8) are available.

## Author Contributions

The manuscript was written through contributions of all authors. All authors have given approval to the final version of the manuscript.

## Funding Sources

This research was funded by the Russian Science Foundation, grant number 18-74-00092.

## Abbreviations

FAD,: flavin adenine dinucleotide;
FMN,: flavin mononucleotide;
FP,: fluorescent protein;
LOV domain,: light oxygen voltage domain;
PDB,: Protein Data Bank;
WT,: wild type.

## Acknowledgements

The diffraction data were collected on the beamlines ID23-1, ID29 and ID30B at the European Synchrotron Radiation Facility (ESRF), Grenoble, France. We are grateful to local contacts at ESRF for providing assistance in using these beamlines, to Daria Melikhova for help with expression of the Q148R variant, and to Ulrich Krauss for fruitful discussions. This research was funded by the Russian Science Foundation, grant number 18-74-00092.

## Notes

### Competing Interest Statement

The authors have declared no competing interest.

