## Supporting information for "Insights into the mechanisms of LOV domain color tuning from a set of high-resolution X-ray structures"

Supporting Table S1

Supporting Figures S1-S8

**Table S1.** Crystallographic data collection and refinement statistics.

| Variant<br>PDB ID | Q148N<br>6YWG | Q148D<br>6YWH | Q148E<br>6YWI | Q148H<br>6YWQ | Q148H<br>6YWR | Q148K<br>6YX4 | Q148K<br>6YX6 | Q148K<br>6YXB | Q148R<br>6YXC |
|---|---|---|---|---|---|---|---|---|---|
| **Data collection** | | | | | | | | | |
| Space group | $P2_12_12$ | $P2_12_12$ | $P2_12_12$ | $P2_12_12$ | C2 | $P2_12_12$ | $P2_12_12$ | $P2_1$ | $P2_12_12$ |
| Cell dimensions | | | | | | | | | |
| *a*, *b*, *c* (Å) | 53.72, 110.56, 38.99 | 53.66, 110.47, 39.04 | 53.55, 109.84, 38.95 | 53.73, 110.35, 38.87 | 110.97, 54.03, 39.23 | 57.66, 110.56, 39.01 | 57.77, 110.84, 39.20 | 39.10, 110.54, 56.94 | 57.42, 110.17, 38.96 |
| *α*, *β*, *γ* (°) | 90, 90, 90 | 90, 90, 90 | 90, 90, 90 | 90, 90, 90 | 90, 98.5, 90 | 90, 90, 90 | 90, 90, 90 | 90, 91.07, 90 | 90, 90, 90 |
| Resolution (Å) | 110.56-1.45 (1.48-1.45) | 55.24-1.07 (1.09-1.07) | 54.92-1.13 (1.15-1.13) | 53.73-1.27 (1.29-1.27) | 54.87-1.50 (1.53-1.50) | 39.90-1.36 (1.38-1.36) | 39.99-1.50 (1.54-1.50) | 110.54-1.50 (1.53-1.50) | 39.75-1.63 (1.76-1.63) |
| $\langle I/\sigma I \rangle$ | 18.5 (2.0) | 16.1 (2.0) | 12.3 (2.4) | 14.3 (2.3) | 9.5 (2.3) | 15.3 (2.0) | 14.8 (2.9) | 10.6 (2.6) | 16.7 (2.4) |
| *CC1/2 (%)* | 99.9 (79.1) | 99.9 (66.5) | 99.1 (80.1) | 99.7 (64.3) | 99.5 (75.3) | 99.9 (79.9) | 99.9 (88.1) | 99.7 (67.6) | 100.0 (74.6) |
| Completeness (%) | 98.6 (99.9) | 98.4 (98.3) | 99.1 (98.5) | 99.9 (99.8) | 96.4 (99.2) | 97.6 (94.4) | 86.8 (59.9) | 98.9 (98.1) | 71.7 (18.0) |
| Ellipsoidal completeness** (%) | | | | | | | 96.2 (98.3) | | 90.9 (53.6) |
| Multiplicity | 12.9 (13.3) | 6.4 (6.0) | 6.3 (5.9) | 6.3 (5.5) | 3.2 (3.2) | 5.9 (5.8) | 5.6 (5.6) | 3.0 (3.0) | 6.4 (6.5) |
| Unique reflections | 41453 (2101) | 101277 (4970) | 86013 (4192) | 61883 (3005) | 35488 (1785) | 53232 (2525) | 35712 (1786) | 76240 (3706) | 22625 (1131) |
| **Refinement** | | | | | | | | | |
| Resolution (Å) | 55.28-1.45 | 48.31-1.07 | 48.18-1.13 | 38.90-1.27 | 54.87-1.50 | 39.94-1.36 | 39.99-1.50 | 56.95-1.50 | 39.75-1.65 |
| No. reflections | 39327 | 96086 | 81699 | 58718 | 33688 | 50560 | 34107 | 72508 | 20790 |
| $R_{work}/R_{free}$ (%) | 17.8/19.9 | 17.6/18.8 | 17.2/18.1 | 16.1/19.1 | 17.6/20.0 | 14.0/17.2 | 17.4/20.2 | 18.4/20.3 | 18.2/21.2 |
| No. atoms | | | | | | | | | |
| Protein | 1740 | 1719 | 1727 | 1732 | 1726 | 1615 | 1645 | 3253 | 1639 |
| FMN | 62 | 62 | 62 | 62 | 62 | 62 | 62 | 124 | 62 |
| Water and others | 223 | 246 | 235 | 198 | 196 | 340 | 361 | 496 | 214 |
| Average *B* factors (Å$^2$) | | | | | | | | | |
| Protein | 16.7 | 13.4 | 15.6 | 15.5 | 17.4 | 21.0 | 14.3 | 13.2 | 27.2 |
| FMN | 12.5 | 9.6 | 11.0 | 11.2 | 14.6 | 15.4 | 8.1 | 8.4 | 19.3 |
| Water and others | 27.6 | 26.2 | 25.4 | 26.8 | 29.5 | 34.6 | 30.2 | 25.9 | 36.0 |
| R.m.s. deviations | | | | | | | | | |
| Protein bond lengths (Å) | 0.005 | 0.008 | 0.008 | 0.011 | 0.005 | 0.007 | 0.005 | 0.005 | 0.003 |
| Protein bond angles (°) | 1.3 | 1.4 | 1.4 | 1.7 | 1.7 | 1.4 | 1.3 | 1.3 | 1.2 |
| Ramachandran analysis | | | | | | | | | |
| Favored (%) | 99 | 100 | 100 | 99 | 99 | 97 | 97 | 98 | 97 |
| Outliers (%) | 0 | 0 | 0 | 0 | 0 | 0 | 0.5 | 0 | 0 |

* The data for the lowest and highest resolution shells are shown in parentheses. R.m.s.: root mean square.

** Ellipsoidal completeness is reported for the cases where the data was anisotropically truncated and corrected

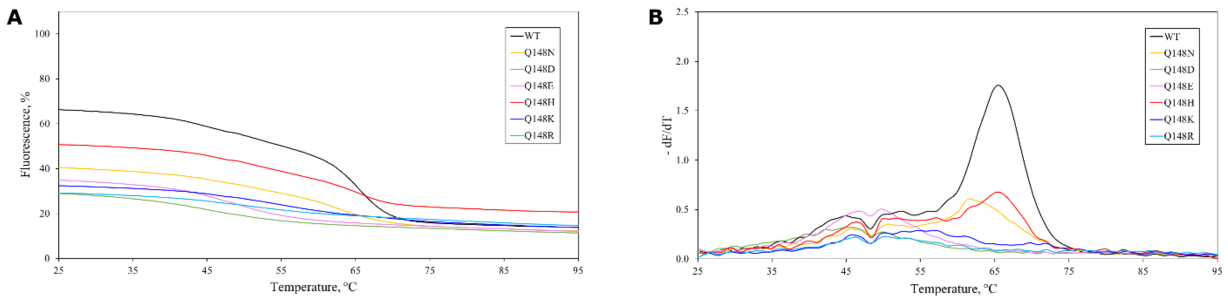


**Figure S1.** Dependence of CagFbFP variants' fluorescence on temperature during cooling. A) Temperature-induced recovery of fluorescence. B) Derivatives of fluorescence traces. Each experiment was conducted independently for five times, and the data were averaged for plotting. Characteristic refolding temperatures are summarized in Table 1.

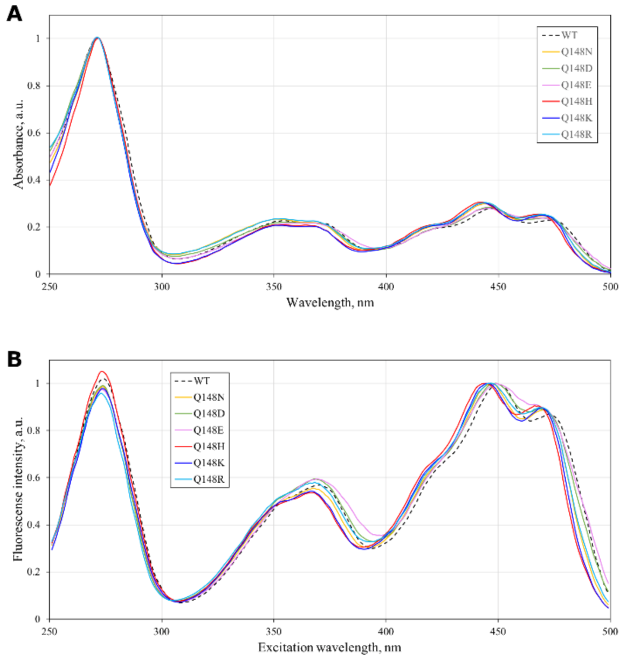


**Figure S2.** Spectroscopic properties of CagFbFP variants at pH 8 in the wavelength range 250-500 nm. A) Absorption spectra. B) Fluorescence excitation spectra.

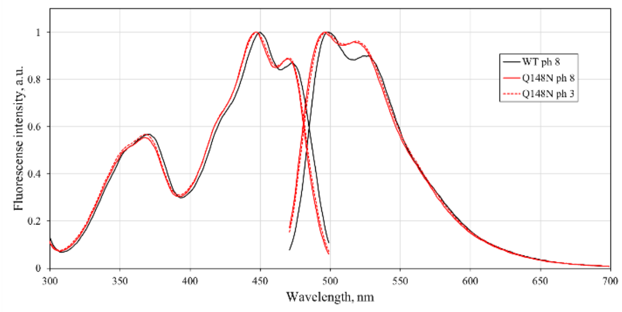


**Figure S3.** Dependence of CagFbFP Q148N fluorescence excitation and emission spectra on pH.

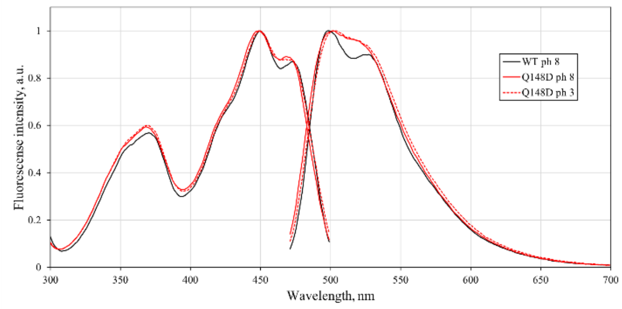


**Figure S4.** Dependence of CagFbFP Q148D fluorescence excitation and emission spectra on pH.

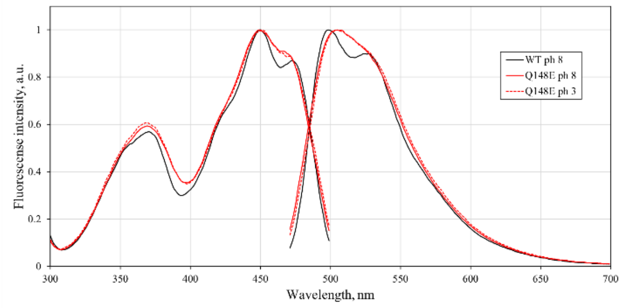


**Figure S5.** Dependence of CagFbFP Q148E fluorescence excitation and emission spectra on pH.

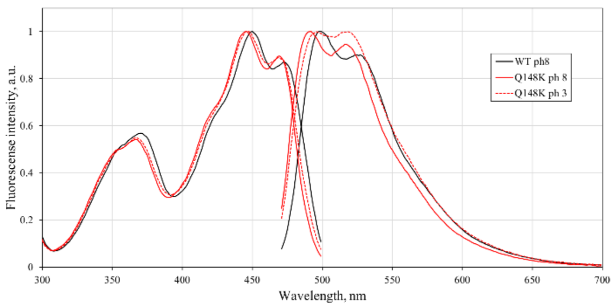


**Figure S6.** Dependence of CagFbFP Q148K fluorescence excitation and emission spectra on pH.

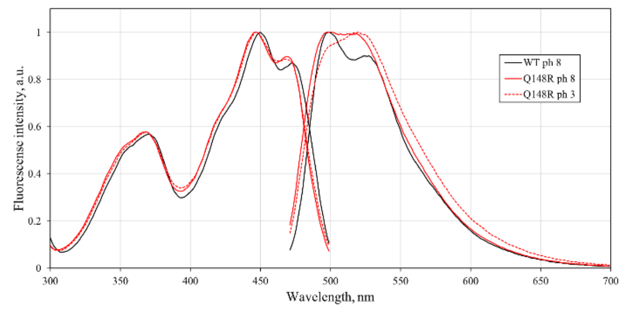


**Figure S7.** Dependence of CagFbFP Q148R fluorescence excitation and emission spectra on pH.

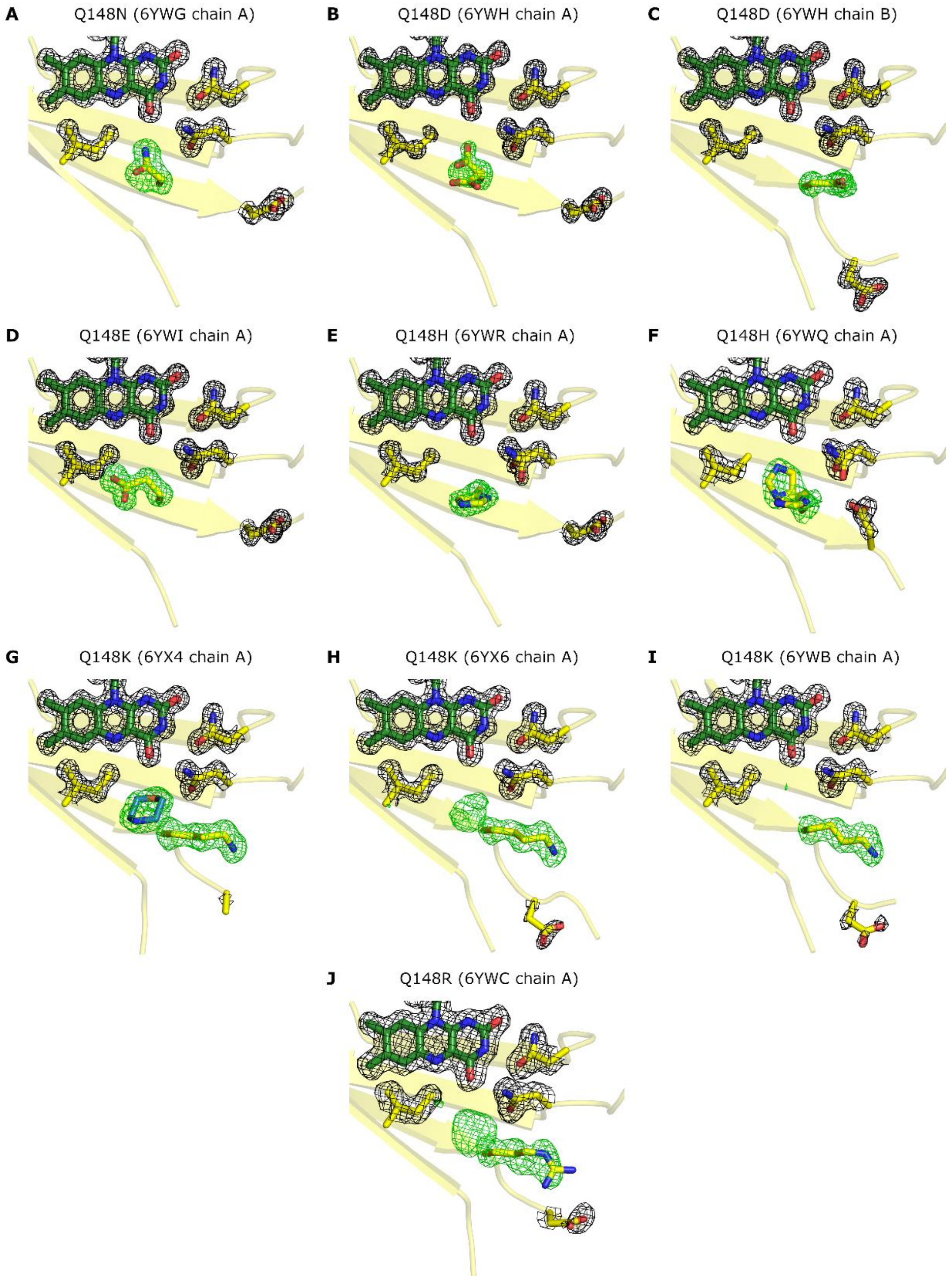


**Figure S8.** Exemplary $2F_o$-$F_c$ (black) and omit (polder, green) maps for Q148X variants contoured at the levels of 1.5 and 4 × r.m.s., respectively. In panels F and J, the polder maps are contoured at 3 × r.m.s., and in panels H and I at 3.5 × r.m.s. for clarity.
